# Intraflagellar transport protein 74 is essential for mouse spermatogenesis and male fertility by regulating axonemal microtubule assembly in mice

**DOI:** 10.1101/457804

**Authors:** Shi Lin, Zhou Ting, Huang Qian, Zhang Shiyang, Li Wei, Zhang Ling, Hess Rex A, Pazour Gregory J, Zhang Zhibing

## Abstract

IFT74 is a component of the core intraflagellar transport (IFT) complex, a bidirectional movement of large particles along the axoneme microtubules for cilia formation. In this study, we investigated its role in sperm flagella formation and discovered that mice deficiency in IFT74 in male germ cells were infertile associated with low sperm counts and immotile sperm. The few developed spermatozoa displayed misshaped heads and short tails. Transmission electron microscopy revealed abnormal flagellar axoneme in the seminiferous tubules where sperm are made. Clusters of unassembled microtubules were present in the spermatids. Testicular expression levels of IFT27, IFT57, IFT81, IFT88 and IFT140 were significantly reduced in the mutant mice, with the exception of IFT20 and IFT25. The levels of ODF2 and SPAG16L proteins were also not changed. However, the processed AKAP4 protein, a major component of the fibrous sheath, a unique structure of sperm tail, was significantly reduced. Our study demonstrates that IFT74 is essential for mouse sperm formation, probably through assembly of the core axoneme and fibrous sheath, and highlights a potential genetic factor (IFT74) that contributes to human infertility in men.

## Introduction

Cilia and flagella are microtubule-based organelles that are found on the surface of most eukaryotic cells. They have been adapted for a variety of functions such as cellular motility, directional fluid movement, cellular signaling and sensory reception (Wheatley, 1995; Bray, 2001; Praetorius and Spring, 2005). The assembly and maintenance of cilia and flagella depend on intraflagellar transport (IFT), which is a bidirectional movement of large particles along the microtubule-based axoneme between the cell body and the distal tip of cilia/flagella (Kozminski et al., 1993; Kozminski et al., 1995). The IFT machinery is composed of kinesin-2, cytoplasmic dynein, and protein complexes known as A and B (Cole et al., 1998; Pazour et al., 1999; Piperno and Mead, 1997). The IFT-A complex consists of six subunits (IFT43, IFT121/WDR35, IFT122, IFT139/TTC21B, IFT140, and IFT144/WDR19) and other ancillary proteins, which is powered by dynein 2 to the cilia/flagella tip (Ishikawa and Marshall, 2011; Taschner et al., 2012). The IFT-B complex contains a salt-stable core complex of nine subunits (IFT22, IFT25, IFT27, IFT46, IFT52, IFT70, IFT74, IFT81, and IFT88) and several peripheral components (IFT20, IFT54, IFT57, IFT80, IFT172, and others), which is driven by kinesin-2 to the cell body (Taschner et al., 2012; Kozminski et al., 1993; Cole et al., 1998; Pazour et al., 1999; Porter et al., 1999; Taschner and Lorentzen, 2016). Defects in the assembly and functions of cilia/flagella, including those in IFT machinery, have been associated with an expanding list of human diseases, including polycystic kidney, obesity, respiratory defects, retinal degeneration, brain and skeletal malformation, as well as infertility (Hildebrandt et al., 2011; Brown and Witman, 2014; Fliegauf et al., 2007).

Although the identities of most IFT proteins are known, the specific function of each subunit are poorly understood. As previously reported, the IFT-B complex is much less stable than IFT-A complex in flagellar isolated from *Chlamydomonas* (Signor et al., 1999; Lucker et al., 2005). Within the core subset of IFT-B proteins, IFT25/27, IFT88/52/46, and IFT81/74/72 are considered to interact to form a heterodimer, a ternary complex, and a heterotetramer in a ratio of 2:1:1, respectively (Lucker et al., 2005; Wang et al., 2009; Lucker et al., 2010). It should be noted that the direct interaction of IFT74 and IFT81 through central and C-terminal coiled-coil domains are sufficient to stabilize IFT-B complex (Taschner et al., 2011; Lucker et al., 2005). Bhogaraju and colleagues found that the N-termini of both proteins form a tubulin-binding module that enhances the affinity of this interaction. Meanwhile, the N-terminus of IFT74 interacted with the highly acidic C-terminal tails (called as E-hooks) of β-tubulin to enhance the binding affinity of *Homo sapiens* IFT81N bound tubulin by ̴18 fold (Bhogaraju et al., 2013). The transport of tubulin to the tip of cilia is not only crucial for cilium assembly but is also essential for maintenance (Marshall and Rosenbaum, 2001; Stephens, 1997). It is suggested that IFT74/81, especially IFT74, plays a critical role in binding and transport of tubulin to the tip of the cilium and the extent of ciliogenesis. In addition, the region near the N-terminus of IFT74 coiled-coil domain 1 is particularly required for the normal association of IFT-A with IFT-B in the cell body of flagella and IFT injection frequency (Brown et al., 2015). Thus, IFT74 combined with IFT81 has an important impact on IFT-B complex stabilization, tubulin transport, and cilium formation and length.

It is demonstrated that the mammalian homologue of the IFT74/72 (referred to as IFT71 in the first report) protein of *Chlamydomonas* is *capillary morphogenesis gene (CMG)-1* (Iomini et al., 2004; Masuda et al., 1997; Bell et al., 2001). Fujino et al revealed that the mouse *CMG-1* gene was specifically expressed in male germ-line stem cells but not in embryonic stem cells (Fujino et al., 2006). As previously reported, CMG-1 is localized in the primary cilia and centrosomes, but not in the nucleus of human umbilical vein endothelial cells (HUVEC) (Iomini et al., 2004). However, Ohbayashi and colleagues have recently identified that CMG-1 is broadly distributed in both the cytoplasm and nucleus of GC-2 cells, a mouse pre-meiotic spermatocyte-derived cell line. Moreover, CMG-1 is required for cell division and niche interactions in the early stages of spermatogenesis in the testis (Ichiro Ohbayashi et al., 2010).

Even though IFT74 is indispensable for the proliferation of male germ-line stem cells in the mouse testis, the physiological roles of IFT74 in the process of spermatogenesis remain largely unknown. Thus, conditional knockout strategy was utilized to investigate the potential role of IFT74 in sperm flagella formation and male fertility. With this approach, our laboratory has disrupted mouse *Ift20*, *Ift25* and *Ift27* genes in male germ cells in mice, and we found that all of them were required for male fertility, spermiogenesis and sperm flagella formation. The study presented here generated the male germ cell-specific *Ift74* knockout mice by breeding floxed *Ift74* mice with *Stra8-iCre* mice as previous studies reported. We discovered that the conditional *Ift74* knockout mice did not show any gross abnormalities, but complete male infertility with dramatically decreased sperm counts and aberrant structures of sperm heads and flagella were present. In the conditional *Ift74* knockout mice, expressions of IFT81 protein, an important IFT74 binding partner, and components of other IFT complex such as IFT27, IFT57, IFT88 and IFT140 proteins were significantly reduced. Our findings suggested that IFT74 is essential for normal mouse spermatogenesis, sperm flagella formation and male fertility in mice.

## Results

### Mouse IFT 74 protein expression and localization in mice

IFT74 protein expression was examined in multiple mouse tissues including heart, brain, spleen, lung, liver, kidney, muscle and testis by Western blot analysis using a highly sensitive Femto system. While the IFT74 protein was present in the other organs containing cilia-bearing cells, such as brain, lung, and kidney, it was highly expressed in the testis (Fig. 1A). The level of IFT74 protein was subsequently evaluated in mouse testis at different times during the first wave of spermatogenesis. The IFT74 protein was detectable beginning day 12, which exhibited a significant increase in abundance from day 20 and after (Fig. 1B).

**Fig 1.**
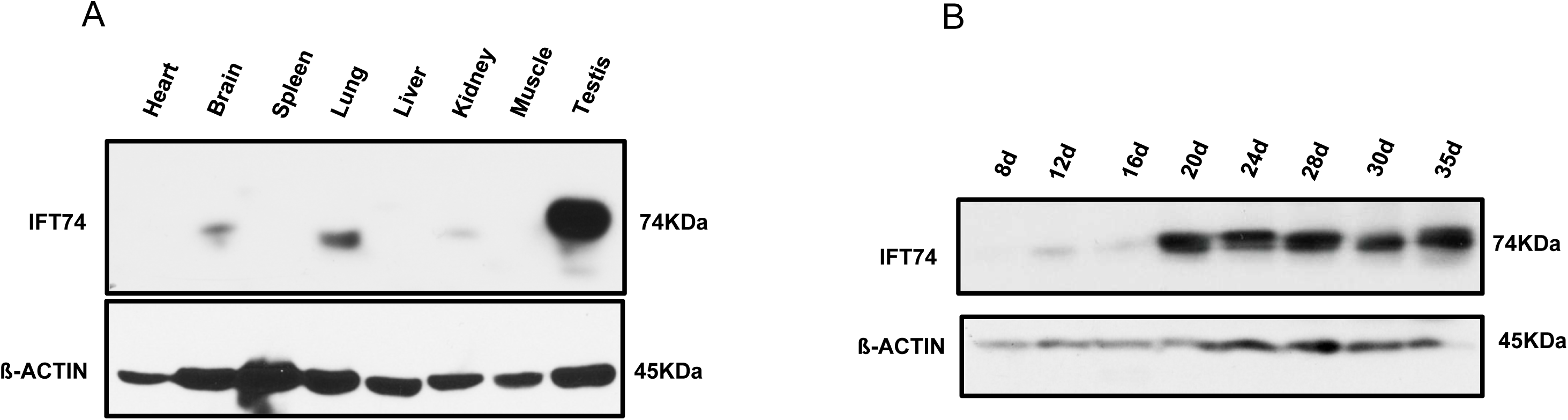
Mouse IFT74 protein is highly expressed in the testis and developmentally regulated during spermatogenesis. A. Western blot analysis of mouse IFT74 protein, using a high sensitive Femto system. Notice that IFT74 is highly expressed in the testis and is also present in the tissues bearing motile and primary cilia, including brain, lung and kidney. B. IFT74 expression during the first wave of spermatogenesis. A representative Western blot result shows that its expression is significantly increased at day 20 after birth.

In addition, immunofluorescence staining was conducted to investigate where the IFT74 protein was localized in isolated germ cells from testis in wild-type mice. Non-specific staining was not observed in cells where no antibody was added (Fig. 2Aa). In control mice, the specific IFT74 signal was strongly expressed not only in the vesicles of spermatocytes and round spermatids (Fig. 2Ab, c), but appeared also in the acrosome and centrosome regions of elongating spermatids (Fig. 2Ad; Fig. 2Bb) and in developing sperm tails (Fig. 2Ae). The cells were further double stained with an acrosome marker, the lectin peanut agglutinin, and IFT74 partially co-localized over the acrosomal region (Fig. 2Ba, b).

**Fig 2.**
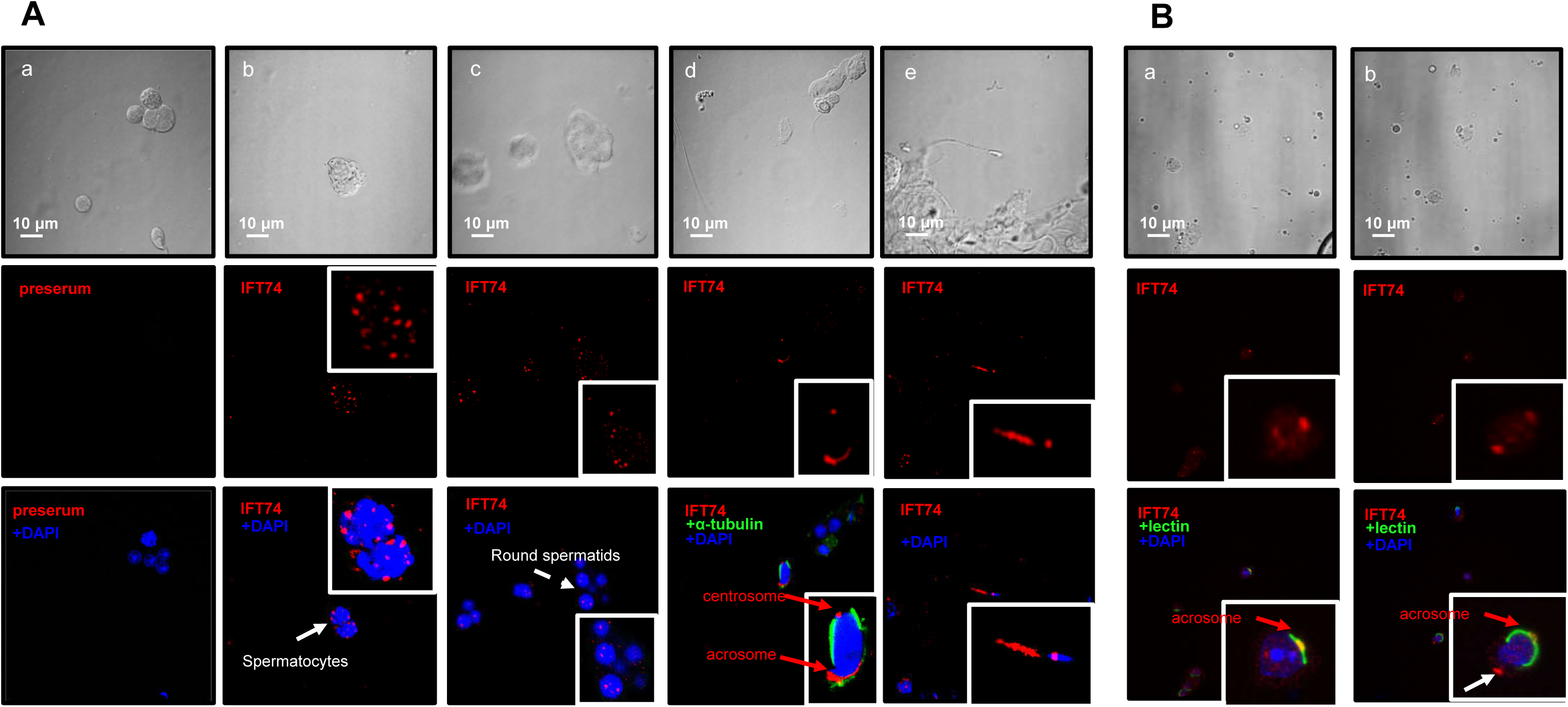
Localization of IFT74 in male germ cells. A. Immunofluorescence staining of IFT74 in germ cells of wild type mice. The top row show the germ cells with phase contrast microscopy. No specific signal was detected in the negative control using preimmune serum (a). It is present as vesicles in spermatocytes (arrow and insert in b) and round spermatids (dashed arrow and insert in (c). It appears to be present in the acrosome and centrosome of elongating spermatid (d), and developing tail (white arrowhead in e); B. The cells were double stained with a lectin acrosome marker, peanut agglutinin. IFT74 was co-localized with lectin. In addition, it was also present as a dot at the opposite region of the acrosome (white arrow), presumably the centriole.

### Generation of conditional *Ift74* knockout mice

The expression pattern and localization of IFT74 suggested an essential role for the protein in spermatogenesis; therefore, male germ cell-specific conditional knockout mice were generated by crossing *Ift74^flox/flox^* females with *Stra8-iCre* transgenic mice (Fig S1). Testicular IFT74 protein expression in control and conditional *Ift74* mutant mice was determined by Western blot analysis and immunofluorescence staining. Western blot results showed that IFT74 protein was nearly absent in the testes of the homozygous mutant mice, whereas it was robustly expressed in all control mice (Fig. 3A). However, specific IFT74 signal by immunofluorescence staining was absent in the knockout mice (not shown).

**Fig 3.**
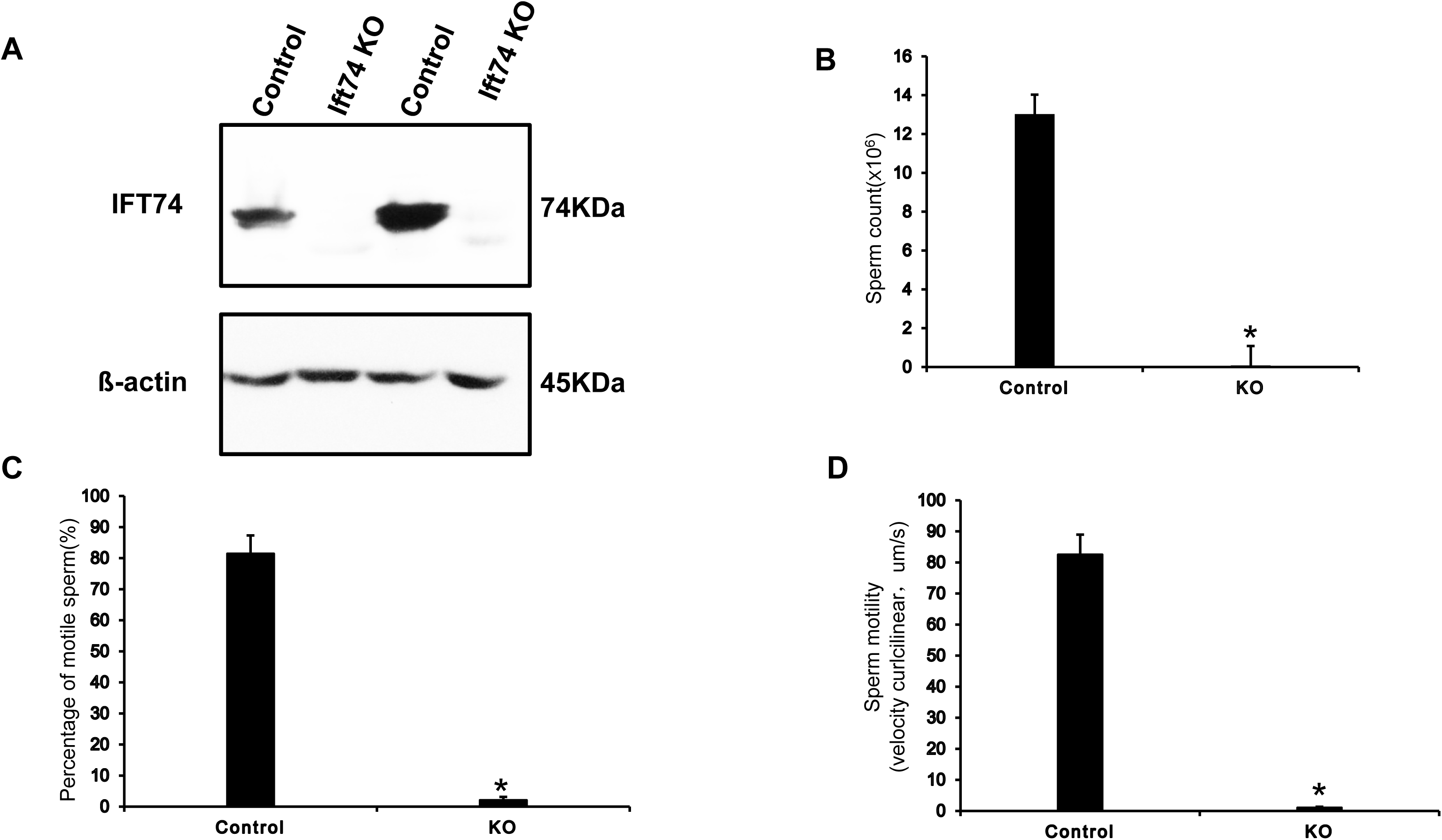
Significant reduction in sperm numbers and motility in the conditional *Ift74* knockout mice. A. Western blot analysis of testicular IFT74 protein expression in control and conditional *Ift74* knockout mice. Notice that IFT74 protein was missing in the knockout mice. Epididymal sperm were collected and physiologic parameters were compared between the control and *Ift74* knockout mice. In *Ift74* knockout mice, there was a significant reduction (* p< 0.05) in sperm counts (B), percentage of motile sperm (C), and sperm motility (D).

### Homozygous conditional *Ift74* KO males were infertile, with significantly reduced sperm counts and motility

All mutant mice survived to adulthood and did not show gross abnormalities. To evaluate fertility, 2-3 months old controls and homozygous *Ift74* KO males were bred to 2-3 months old wild-type females for at least two months. All six control males were fertility and sired normal size litters. However, all six *Ift74* mutant males were infertile and did not produce any litters during the breeding period (Table 1). To investigate the mechanisms that underline the infertility, sperm numbers and motility were examined. Sperm counts were reduced in the conditional *Ift74* knockout mice (Fig. 3B) and there was a significant reduction in the percentage of motile sperm (Fig. 3C). Sperm motility was also dramatically reduced in the conditional *Ift74* knockout mice (Fig. 3D, supplemental movies).

**Table 1.** Homozygous *Ift74* knockout males were infertile.

| Genotype | Fertility | Litter size<br>(n=6) | Testis/body<br>weight (n=6, mg/g) |
| --- | --- | --- | --- |
| Control | 6/6 | 7±2 | 9.47±0.68 |
| KO | 0/6 | 0 | 8.06±0.40 |

To test fertility, 2-3 months old control and conditional *Ifi74* knockout mice were bred to 2-3 months old wild-type females for at least 2 months. Litter size was recorded for each mating.

### Abnormal epididymal sperm in the conditional *Ift74* KO mice

To further identify sperm changes associated with the infertility phenotype, morphology of cauda epididymal sperm was analyzed by light microscopy (Fig S2) and SEM (Fig. 4, Fig S3). Sperm from control mice had well-shaped heads and long, smooth tails. However, only few sperm were recovered sperm from cauda epididymides from *Ift74* knockout mice and all were abnormal. A variety of abnormalities were observed in the mutant sperm, including very short tails and mostly abnormally shaped heads, with only rare sperm having a normal appearing head.

**Fig 4.**
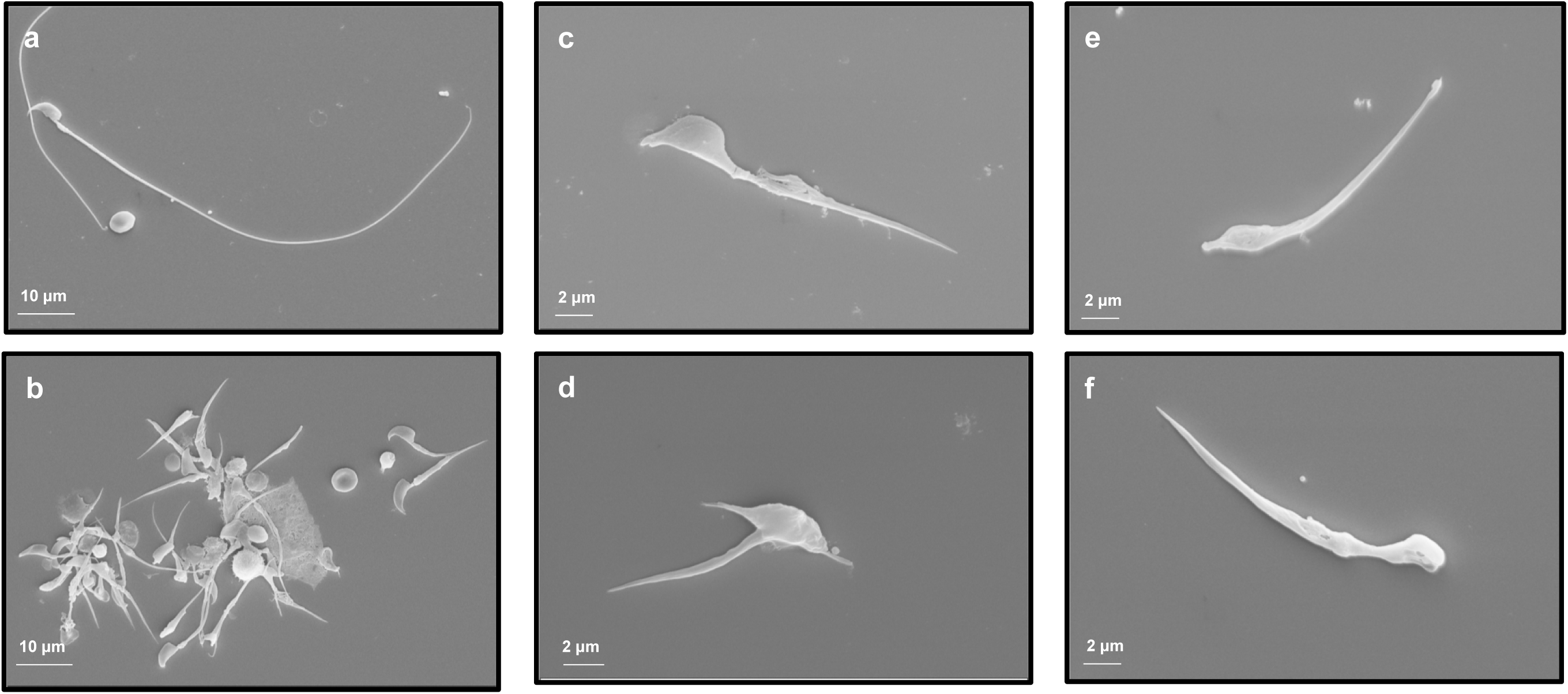
Abnormal epididymal sperm in the conditional *Ift74* knockout mice. Examination of epididymal sperm by SEM. a: Representative image of epididymis sperm with normal morphology from a control mouse. The sperm has a normally shaped head and a long, smooth tail; “b” to “d”: Representative images of epididymal sperm from a conditional *Ift74* knockout mice. All sperm have short tails and most have grossly abnormal heads. “c” to “f” show sperm with both abnormal heads and a short abnormal tails.

### Abnormal spermiogenesis in the conditional *Ift74* KO mice

To analyze changes in spermatogenesis that may contribute to low sperm counts and abnormal sperm morphology, histology of testes in control and *Ift74* KO mice were examined (Fig. 5A, Fig S4A). There was no significant difference in testis/body weights between controls and mutant mice (Table 1). In control mice, testes exhibited an integrated and normal spermiogenesis (Fig. 5Aa). However, testes of *Ift74* KO mice showed a failure of spermiation in stage VIII, with abnormal step 16 spermatids (Ab16) being phagocytized, although in the same tubule step 8 round spermatids and pachytene spermatocytes (P) appeared normal. Also, normal residual bodies were not formed. Instead, small pieces of germ cell cytoplasm (Cy) were retained at the luminal border (Fig. 5Ab) or sloughed into the lumen, as found in the epididymis (Fig. 5Bb). In mutant stage XI, abnormal step 11 spermatids (Ab 11) were observed with misshaped heads and the absence of sperm tails (Fig. 5Ac). Mutant stage I tubules showed normal round spermatids, but there were abnormal step 13 elongating spermatids (Ab13) lacking tails. Excess cytoplasm (Cy) of the elongating spermatids appeared to be sloughed into the lumen (Fig.5Ad).

**Fig 5.**
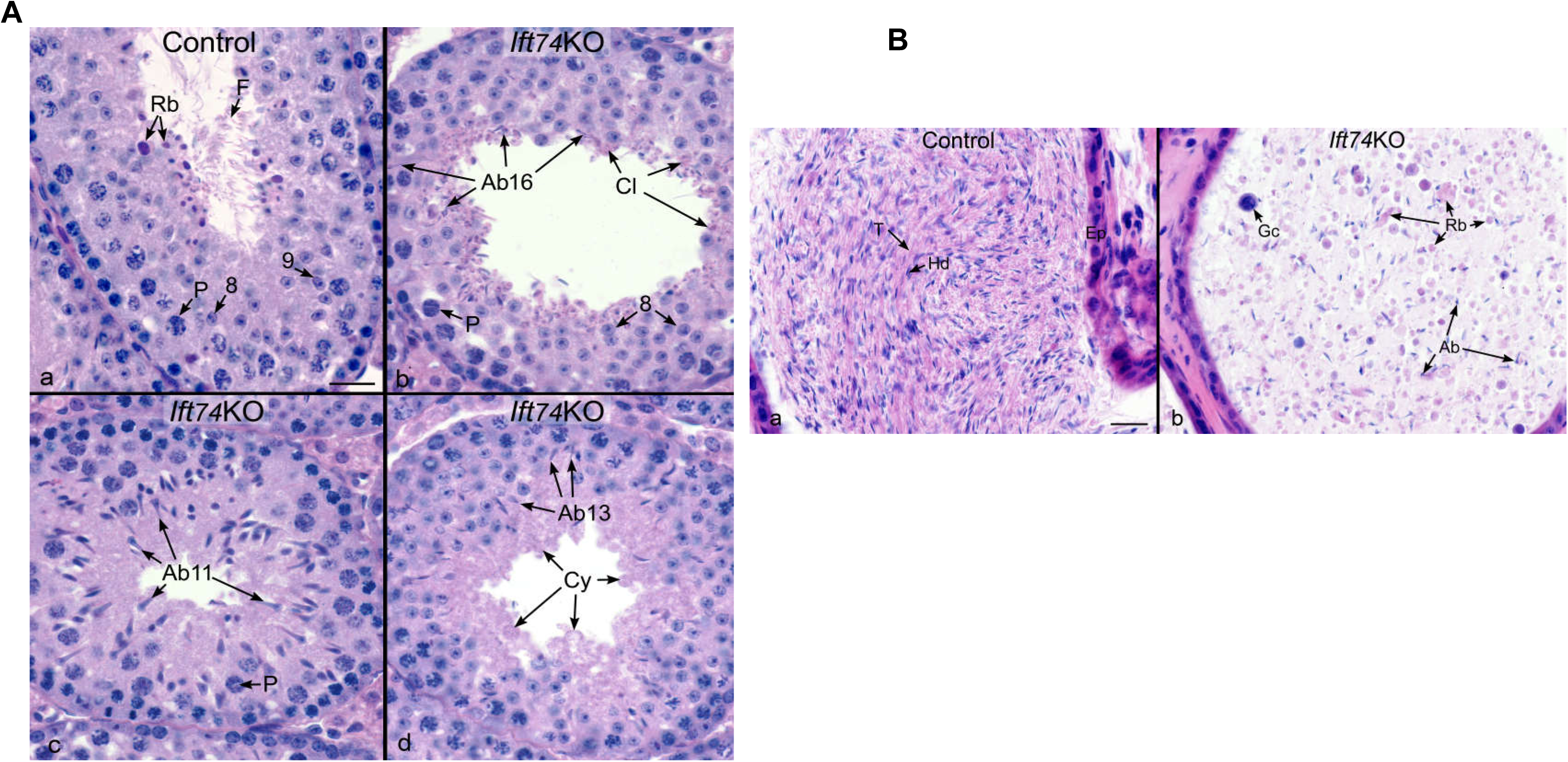
Abnormal spermiogenesis in the conditional *Ift74* knockout mice. A. Testis histology from control and *Ift74* KO mice showing cross sections of seminiferous tubules. Bar = 20μm. a) Control seminiferous tubule Stage VIII-IX, showing steps 8 and 9 round spermatids, flagella (F) of sperm being released into the lumen and residual bodies (Rb) of germ cell cytoplasm being phagocytized by Sertoli cells. P, pachytene spermatocyte. b) *Ift74* KO seminiferous tubule in Stage VIII, showing normal step 8 round spermatids and pachytene spermatocytes (P). Abnormal step 16 spermatids (Ab16) are seen being phagocytized and failing to spermiation. Residual bodies are not forming but small pieces of germ cell cytoplasm (Cy) are retained at the luminal border. c) *Ift74* KO Stage XI with abnormal step 11 spermatids (Ab11) with abnormally shaped heads and absence of tails. P, pachytene spermatocyte. d) *Ift74* KO tubule showing normal round spermatids but abnormal step 13 elongating spermatids (Ab13) that are lacking tails. Excess cytoplasm (Cy) of the elongating spermatids appears to be sloughed into the lumen. B. Cauda epididymis from control and *Ift74* KO mice. Bar = 20 μm. a) Control epididymis showing an epithelium (Ep) lining the lumen that is filled with normal sperm aligned with their heads (Hd) and tails (T). b) *Ift74* KO epididymis showing a lumen filled with numerous, large cytoplasmic bodies that are likely residual bodies (Rb) and sperm with abnormal heads (Ab) and short or absent tails. Gc, sloughed round spermatid.

Abnormal spermatogenesis in the *Ift74* knockout mice was also manifested by the presence of abnormal luminal contents in the cauda epididymis (Fig. 5B, Fig S4B). In control mice, the lumen that was filled with compacted epididymal sperm that were aligned with normal heads and tails (Fig. 5B, left; Fig S4B upper). However, in *Ift74* KO the epididymal lumen contained massive amounts of large abnormal cytoplasmic bodies, residual bodies, sloughed round cells, and abnormal spermatids with short or absent tails (Fig. 5B, right; Fig, S4B lower).

Ultrastructure of seminiferous tubules was determined by TEM in the control and conditional *Ift74* KO mice (Fig. 6, Fig S5). In control mice, a large number of axonemes were present in the lumen (Fig. 6a). However, in the seminiferous tubules of the conditional *Ift74* KO mice, a variety of axonemal abnormalities were discovered. Spermatid abnormalities included the following: a complete absence of the axoneme, disorganized microtubules, clusters of microtubules, and abnormally formed axonemes without the central microtubules. Although the acrosome of spermatids appeared to be normal, there were some abnormally-shaped heads on elongating spermatids (Fig. 6b-h; Fig S5).

**Fig 6.**
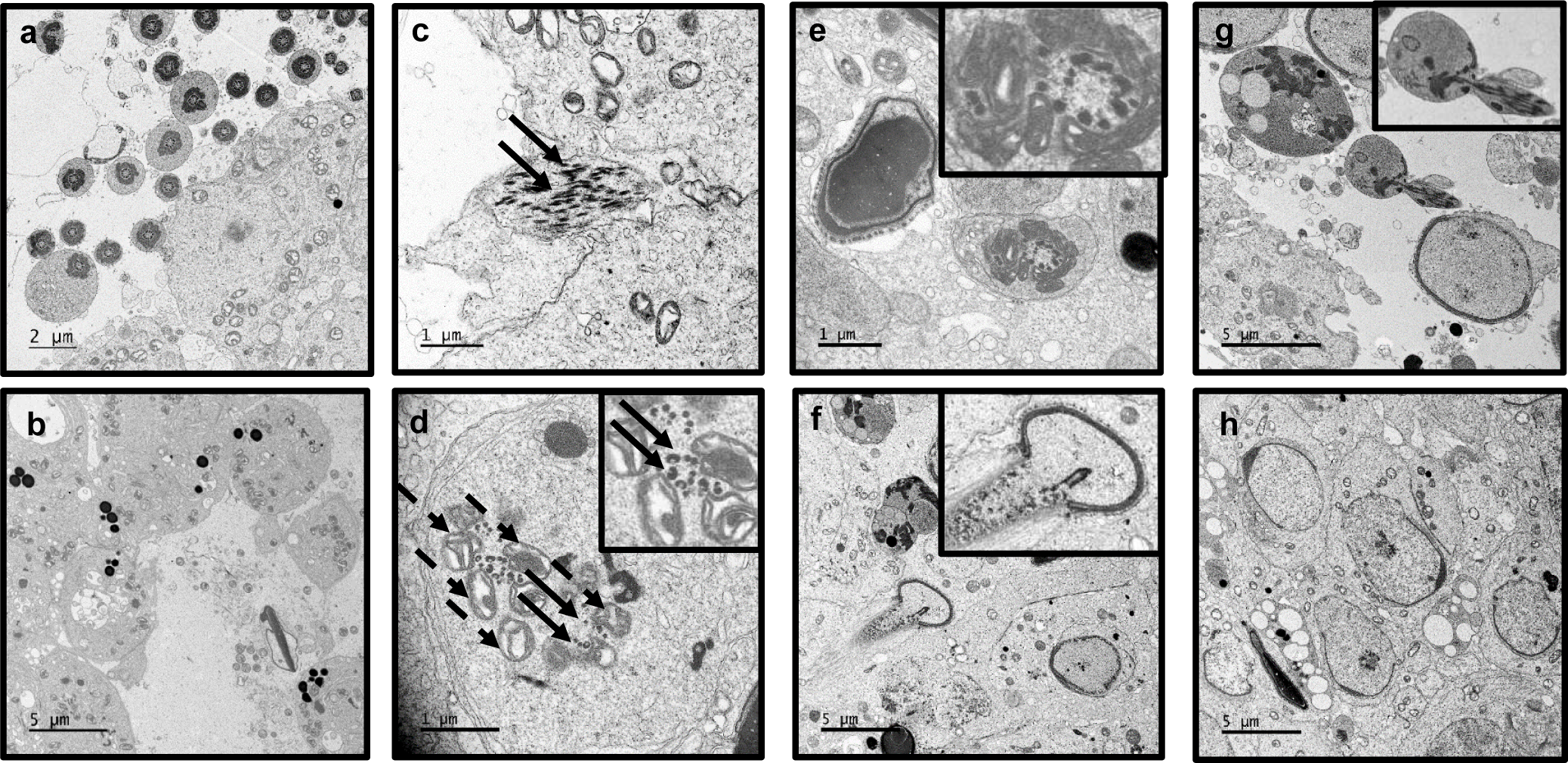
Ultrastructural changes in the testis of conditional *Ift74* knockout mice. Testicular ultrastructural was analyzed in the conditional *Ift74* knockout mice. “a” Control mouse TEM image. Numerous axonemes of sperm tails are seen in the lumen. “b-h” *Ift74* mutant mouse testis. “b” shows the lumen area with the absence of normal axonemal structures. The arrow in “c” points to disorganized microtubules; the insert in “d” shows clusters of microtubules (arrows) and mitochondria (dashed arrows) without a core axoneme structure; the insert in “e” shows an abnormally formed axoneme without the central microtubules; the insert in “f” shows an abnormally formed elongating spermatid; “g” shows sloughed residual bodies and abnormal spermatids; the insert in “g” shows an abnormal spermatid. The developing acrosome appears to be normal (h).

### IFT74 regulates cellular levels of other IFT proteins and some flagellar proteins

To understand how the loss of IFT74 affects other IFT and flagellar proteins, we determined the levels of selective IFT and flagellar proteins in the testis of control and conditional *Ift74* KO mice by Western blot. The protein levels of components of the IFT-B complex, such as IFT27, IFT57, IFT81, IFT88 and IFT140 were significantly decreased in *Ift74* KO mice; however, the IFT20 and IFT25 proteins, two additional components of the IFT-B complex, did not differ from those expressed in control mice (Fig. 7A). Moreover, IFT74 did not affect testicular expression levels of ODF2, a major component of the sperm tail outer dense fibers, and an axonemal central apparatus protein, SPAG16L. However, AKAP4, a processed form of the major component of the fibrous sheath, was significantly reduced in the knockout mice (Fig. 7B).

**Fig 7.**
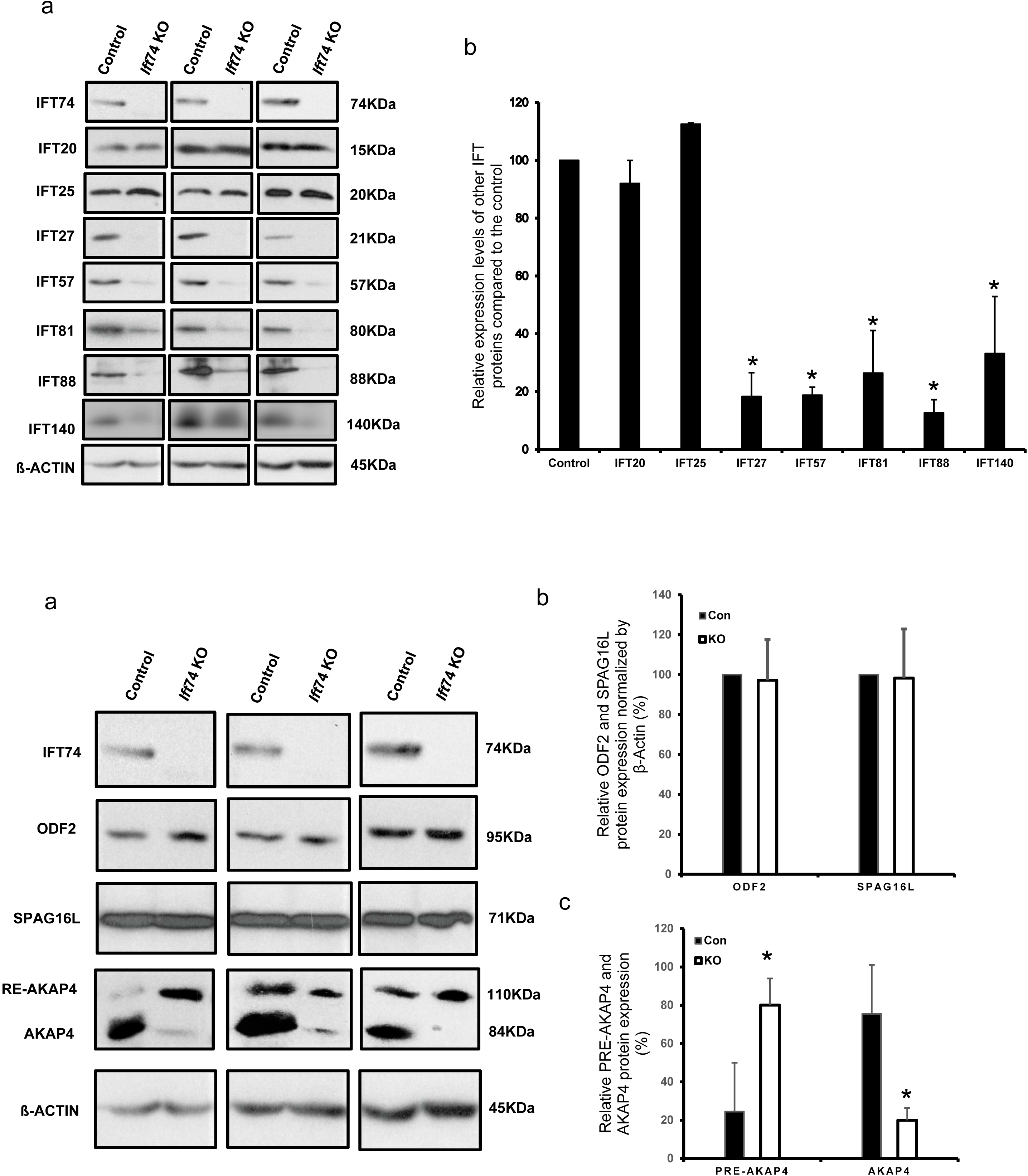
IFT74 regulates expression levels of some IFT and flagellar proteins. A.Examination of selective IFT protein expression levels in the control and conditional Ift74 knockout mice by Western blot. Compared to the controls, the expression levels of IFT27, IFT57, IFT81, IFT88 and IFT140, but not IFT20 and IFT25 are reduced in the conditional *Ift74* knockout mice. a: representative Western blot result; b: quantitative analysis of selective IFT protein expression. B. Examination of testicular expression levels of ODF2, a component of sperm tail outer dense fibers, AKAP4, a major component of fibrous sheath, and SPAG16L, a component of axonemal central apparatus protein in the control and conditional *Ift74* knockout mice. There was no difference in ODF2 and SPAG16L expression. However, the processed AKAP4 was significantly reduced in the conditional *Ift74* knockout mice. β-ACTIN was used as controls. a: representative Western blot result; b: quantitative analysis of ODF2 and SPAG16L; c: quantitative analysis of AKAP4.

## Discussion

In this study, we first characterized the expression pattern of IFT74 in male germ cells, and then examined the reproductive phenotype of conditional *ift74* knockout mice. IFT74 was found to be highly expressed during spermiogenesis, suggesting a specific role in the morphological changes that germ cell undergo during differentiation, particularly formation of the sperm flagellum. This function is consistent with the conserved role of IFT in cilia formation and is further supported by its localization. During spermatogenesis, IFT74 is associated with vesicles in spermatocytes and round spermatids, with the acrosome and centrosome of elongating spermatids, and in the developing sperm tails. This is similar to previous studies (Iomini et al., 2001) showing IFT74 localization in the proximal region of developing flagella and base of the mature flagellum in wild-type cells (Esparza et al., 2013), which suggests that IF74 may be involved in carrying cargo proteins responsible for formation of the sperm flagellum. This hypothesis is strongly supported by data obtained in the conditional *Ift74* knockout mice. The homozygous males were infertile due to low sperm numbers and significantly reduced sperm motility. Abnormalities in sperm morphology we also observed, such as round and distorted heads, short tails, and a great diversity of axonemal and microtubular abnormalities in spermatids from the *Ift74* KO mice. Given that a major ultrastructural observation was the lack of microtubule assembly in spermatids, it is likely that one of IFT74’s major functions in testis is the transport of β-tubulin for microtubule assembly during formation of the axoneme in differentiating spermatids.

It has been shown that IFT74 is expressed in spermatogonia and, most abundantly, in premeiotic spermatocytes (Ichiro Ohbayashi et al., 2010). It is essential for the expression of *cyclin-D2* in the mouse premeiotic spermatocyte-derived GC-2 cell line (Fujino et al., 2006). Furthermore, siRNA-mediated knockdown of *Ift74* in GC-2 cells resulted in a significant reduction of protein levels of cell-adhesion molecules such as E-cadherin protein that is required for the initial cell division of spermatogonial stem cells (Yamashita et al., 2003). These findings from *in vitro* studies were not consistent with our *in vivo* observations from the conditional *Ift74* knockout mice. Even though IFT74 protein is first detected on day 12 after birth in control testes, the knockout mice phenotype was not observed until the spermiogenesis phase. Thus, our results do not support the idea that IFT74 is essential for mitosis and meiotic development of male germ cells.

The function of IFT74 seems to be conserved between *Chlamydomonas reinhardtii* and mouse. In *Chlamydomonas reinhardtii Ift74* null mutant cells had no cilia or only short cilia (Brown et al., 2015). In *Ift74* knockout mice, a consistent observation was the lack of axoneme formation in seminiferous tubules, and when occasionally present the sperm had very short tails. Moreover, the accumulation of disorganized microtubules in mutant spermatid cytoplasm suggests a failure to incorporate tubulin into the flagellum, which would be consistent with the known function of IFT74 as a tubulin carrier. In *Chlamydomonas*, many *Ift74* mutant flagella showed no IFT movements and severe defects of IFT injection (Brown et al., 2015; Wren et al., 2013; Craft et al., 2015). Even though there is limited visibility of IFT protein in the flagellum, IFT frequency and retrograde velocity are both dramatically reduced in the mutant compare with wild-type cells, suggesting that IFT74 could be a master regulator controlling sperm formation by modulating axoneme and microtubule assembly.

The concept that IFT74 may serve as a master regulator of axoneme assembly is supported by Brown and colleagues’ study that IFT74 is required to stabilize IFT-B and flagella assembly (Brown et al., 2015). It has been shown that IFT74 and IFT81 interact directly through not only central and C-terminal coiled-coil domains, but also the N-termini of both proteins to enhance the IFT-B complex stability (Lucker et al., 2005; Bhogaraju et al., 2013). Defect in IFT74 function is likely to affect the interactions of these proteins and the stability of IFT81 formed IFT-B complex. The levels of both core IFT81 and IFT57 proteins are lower in *Chlamydomonas reinhardtii Ift74* mutants than that in wild-type cells (Brown et al., 2015). This function seems to be conserved also in mouse. Expression levels of most IFT components examined, including IFT27, IFT57, IFT81 and IFT88, IFT140 were reduced in the conditional *Ift74* knockout mice, suggesting that IFT74 is a core IFT component that controls the stability of other IFTs. These proteins might be gradually degraded when IFT-B complex is unable to assemble without expression of the *Ift74* gene. This differs from several other conditional *Ift* knockout mice. In conditional *Ift25, Ift27* and *Ift140* knockout mice, expression levels of most IFT proteins, particularly IFT74, were not changed (Zhang et al., 2017; Liu et al., 2017). It has been demonstrated that both peripheral proteins (IFT20 and IFT25) are localized outside of the core IFT-B subcomplex, and their behaviors are possibly different from other IFT-B proteins (Wang et al., 2009; Pedersen et al., 2005; Richey and Qin, 2012; Iomini et al., 2009), which would explain why there was no change in IFT20 and IFT25 expression levels in the *Ift74* knockout mice. In *Ift25* mutant mice, the IFT20 protein level was significantly reduced, but IFT74 protein was still present similar to that of wild-type mice (Liu et al., 2017). It appears that IFT20 and IFT25 form a sub-complex that does not include IFT74 in male germ cells.

Testicular ODF2 and SPAG16L protein levels were not changed in the conditional Ift74 knockout mice. In *Ift20* KO mice, ODF2 and SPAG16L, two sperm flagella proteins, fail to be incorporated into sperm tails (Zhang et al., 2016); thus, the potential role of IFT74 in the localization of these proteins remains to be determined. Interestingly, expression pattern of a sperm fibrous sheath protein, A-Kinase anchor protein (AKAP4) was changed in the absence of IFT74. *Akap4* gene is translated as a full length 110 kDa precursor (pro-AKAP4). The pro-AKAP4 should be transported from cytoplasm to the fibrous sheath assemble site, presumably by IFT. A 26 kDa peptide is processed out at the fibrous sheath assemble site, and the 84 kDa AKAP4 is incorporated into sperm fibrous sheath (Johnson et al., 1997; Turner et al., 1999). In the wild-type mice testis, the AKAP4 seems to be the predominant form, and the pro-AKAP4 level is significant less than AKAP4. However, in the *Ift74* knockout mice, pro-AKAP4 became the predominant form. The reason might be that the pro-AKAP4 is not transported to the fibrous sheath assemble site due to disrupted IFT, and the precursor is not processed. Much research remains if we are to learn how IFT74 modulates sperm structure.

In conclusion, we explored the role of IFT74 in mouse sperm development and male fertility and the findings support that IFT74 is essential for mouse spermatogenesis and specifically the sperm flagellum via the assembly of microtubules during formation of the axoneme. By analyzing the expression of other IFT components and sperm flagella proteins, we have concluded that IFT74 may function as a core component of the IFT-B complex to modulate its stability and for transporting the fibrous sheath precursor for sperm fibrous sheath formation.

## Material and Methods

### Ethics statement

All animal procedures were approved by Wayne State University Institutional Animal Care and Use Program Advisory Committee (Protocol number: IACUC-18-02-0534) in accordance with federal and local regulations regarding the use of nonprimate vertebrates in scientific research.

### Generation of *Ift74* condition knockout mice

The *Ift74^TmIa^* mice were obtained from the KOMP project at Jackson Laboratory and converted to the *Ift74^flox^* allele by FlpE (Farley et al., 2000). *Stra8-iCre* mice were purchased from Jackson Laboratory (Stock No:008208). Transgenic mouse line Stra8-cre expresses improved Cre recombinase under the control of a 1.4 Kb promoter region of the germ cell-specific stimulated by retinoic acid gene 8 (Stra8) (Sadate-Ngatchou et al., 2008). To generate the germ cell-specific *Ift74* KO mice, the same breeding strategy used to generate germ cell-specific *Ift20, Ift25* and *Ift27* knockout mice was used (Zhang et al., 2016; Liu et al., 2017; Zhang et al., 2017). Briefly, three to four-month old *Stra8-cre* males were crossed with three to four-month old *Ift74^flox/flox^* females to obtain *Stra8-iCre; Ift74^flox/+^* mice. The three to four-month old *Stra8-iCre; Ift74^flox/+^* males were crossed back with three to four-month old *Ift74^nox/flox^* females again, and the *Stra8-iCre;Ift74^flox/fIox^* were considered to be the homozygous knockout mice (KO). *Stra8-iCre; Ift74^nox/^+* mice were used as the controls.

Mice were genotyped by PCR using multiplex PCR mix (Bioline, Cat No. BIO25043). To genotype the offspring, genomic DNA was isolated as described previously (Keady et al., 2012). The presence of the *Stra8-iCre* allele was evaluated as previous study (Sadate-Ngatchou et al., 2008), and *Ift74* genotypes were determined as described as previously. The following primers were used for genotyping: *Stra8-iCre* forward: 5-GTGCAAGCTGAACAA CAGGA-3; *Stra8-iCre* reverse: 5-AGGGACACAGCATTGGAGTC-3, and *Ift74* wild-type primer 1:5-CTGAGTGAAAGTGGAGC-3; primer 2 5-CAAGAAAGCTTGGGTCTAGAT-3; KO primer 3: 5-GAATGCATGTGAAATACATTGTGAA-3; primer 4: 5-GAGAAAAGCAGTAATAGTTCTCATCTCC-3.

### Western blot analysis

All tissue samples from three to four-months old mice were homogenized on ice in lysis buffer [(50 mM Tris-HCl pH 8.0, 170 mM NaCl, 1% NP40, 5 mM EDTA, 1 mM DTT and protease inhibitors (Complete mini; Roche diagnostics GmbH)] using Ultra Turrax. Supernatants were collected after being centrifuged at 13000 rpm, for 10 min at 4°C. Protein concentrations were measured using Bio-Rad DCTM protein assay kit (Bio-Rad) by Lowry assay. Proteins were denatured under 95°C for 10 min, then separated by SDS-PAGE and transferred to polyvinylidene fluoride membranes (Millipore, Billerica, MA, USA). Then the Nonspecific sites were blocked in a Tris-buffered saline solution containing 5% nonfat dry milk powder and 0.05% Tween 20 (TBST) for 1 hour at room temperature, and the membranes were incubated with indicated primary antibodies (IFT25: 1:2000, Cat No: 15732-1-AP, ProteinTech; IFT74: 1:2000, Cat No: AAS27620e from ANTIBODY VERIFY; IFT81: 1:1000, Cat No: 11744-1-AP, ProteinTech; β-actin: 1:2000, Cat No: 4967 S, Cell Signaling Antibodies against IFT20, IFT27, IFT57, IFT88 and IFT140 (1:2000) were from Dr. Pazour’s laboratory (Keady et al., 2012, Pazour et al., 2002, Jonassen et al., 2012); AKAP4: 1:4000, from Dr. George Gerton at University of Pennsylvania; ODF2: 1:800, Cat No: 12058-1-AP, ProteinTech; SPAG16L: 1:1000, generated by Z.Z.’s laboratory) at 4°C overnight. After washing three times with TBST, the membranes were incubated with the secondary antibody conjugated with horseradish peroxidase with a dilution of 1:2000 at room temperature for at least 1 h. After washing with TBST twice and a final washing with TBS, the bound antibodies were detected with Super Signal Chemiluminescent Substrate (Pierce, Rockford, IL, USA).

### Assessment of fertility and fecundity

To test fertility and fecundity, 6-week-old or 3-4 months old conditional Ift74 KO and control males were paired with adult wild-type females (3-4 months old). Mating cages typically consisted of one male and one female. Mating behavior was observed, and the females were checked for the presence of vaginal plugs and pregnancy. Once pregnancy was detected, the females were put into separate cages. Breeding tests for each pair lasted for at least three months. The number of pregnant mice and the number of offspring from each pregnancy were recorded.

### Spermatozoa counting

Sperm cells were collected and relocated in warm PBS from cauda epididymides and fixed with 2% formaldehyde for 10 min at room temperature. Followed by washing with PBS, sperm were suspended into PBS again and counted using a hemocytometer chamber under a light microscope, and sperm number was calculated by standard methods as we used previously (Zhang et al., 2006).

### Spermatozoa motility assay

Sperm were collected from the cauda epididymides in warm PBS. Sperm motility was evaluated using an inverted microscope (Nikon, Tokyo, Japan) on a prewarmed slide with a SANYO (Osaka, Japan) color charge-coupled device, highresolution camera (VCC-3972) and Pinnacle Studio HD (version 14.0) software. Movies were taken at 15 frames/sec. For each sperm sample, ten fields were selected for analysis. Individual spermatozoa were tracked using NIH Image J (National Institutes of Health, Bethesda, MD) and the plug-in MTrackJ. Sperm motility was calculated as curvilinear velocity (VCL), which is equivalent to the curvilinear distance (DCL) traveled by each individual spermatozoon in one second (VCL = DCL/t).

### Histology on tissue sections

Adult mice testes and epididymides were fixed in 4% formaldehyde solution in Phosphate-buffered saline (PBS), paraffin embedded, and sectioned into 5 μm slides. Haematoxylin and eosin staining was conducted using standard procedure. Histology was examined using a BX51 Olympus microscope (Olympus Corp., Melville, NY, Center Valley, PA), and photographs were taken with the ProgRes C14 camera (Jenoptik Laser, Germany).

### Isolation of spermatogenic cells and immunofluorescence analysis

Testis from adult mice were dissected in a 15 mL centrifuge tube with 5 mL DMEM containing 0.5 mg/mL collagenase IV and 1.0 μg/mL DNAse I (Sigma-Aldrich) for 30 min at 32°C and shaken gently. Then released spermatogenic cells were washed one time with PBS after centrifuging for 5 min at 1000 rpm and 4°C, and the supernatant was discarded. Afterwards, the cells were fixed with 5 mL of 4% paraformaldehyde (PFA) containing 0.1 M sucrose and shaken gently for 15 min at room temperature. After washing three times with PBS, the cell pellet was resuspended with 2 mL PBS, loaded onto positively charged slides, and stored in a wet box after the sample on slides air-dried. The spermatogenic cells were permeabilized with 0.1% Triton X-100 (Sigma-Aldrich) for 5 min at 37 °C, washed with PBS three times and blocked with 10% goat serum for 30 min at 37°C. Then cells were washed with PBS three times and incubated overnight with an anti-IFT74 antibody (1:200). The primary antibodies used were the same as those used for Western blot analysis, but the dilutions were 10 times higher. Following the secondary antibody incubation, the slides were washed with PBS three times, mounted using VectaMount with 4’, 6-diamidino-2-phenylindole (DAPI)(Vector Laboratories, Burlingame, CA), and sealed with nail polish. Images were captured by confocal laser-scanning microscopy (Zeiss LSM 700).

### Transmission electron microscopy (TEM)

Testis and epididymal sperm from adult mice were fixed in 3% glutaraldehyde/ 1% paraformaldehyde/0.1 M sodium cacodylate, pH 7.4 at 4°C overnight and processed for electron microscopy as reported (Zhang et al., 2017). Images were taken with a Jeol JEM-1230 transmission electron microscope.

### Scanning electron microscopy (SEM)

For SEM analysis, Mouse epididymal sperm were collected and fixed in the same fixative solution as transmission electron microscopy (TEM). The samples were processed by standard methods (Zhang et al., 2006) and images were taken with a Zeiss EVO 50 XVP SEM at Microscopy Facility, Department of Anatomy and Neurobiology, Virginia Commonwealth University.

### Statistical analysis

Analysis of variance (ANOVA) test was used to determine statistical difference; the 2-tailed student’s t-test was used for the comparison of frequencies. Significance was defined as *P* < 0.05.

## Supporting information

**Fig S1. Representative PCR results showing mice with different genotypes.** Upper panel: primer set to analyze *Ift74* genotyping; lower panel: primer set to detect *Cre.*

**Movies S1. Examples of sperm motility patterns from the control mice and the conditional *Ift74* mutant.** The movies are short segments of freshly isolated, non-capacitated sperm from a control (A) and the conditional *Ift74* mutant mice (B). All segments were recorded with a DAGE-MTI DC-330 3CCD camera and a Canon Optura 40 digital camcorder. Segments were assembled into the video using iMovie HD on a Dual 1GHz 414 PowerPC Processor G4 Apple Macintosh computer. Movie A. A representative movie from a control mouse. Note that most sperm are motile and display vigorous flagellar activity and progressive, long-track forward movement. Movie B. A representative movie from a conditional *Ift74* mutant mouse. Notice that there are fewer sperm compared to the control mice, with the same dilution, and all sperm are immotile. A large number of degenerated cells are also present.

**Fig S2. Morphological Examination of epididymal sperm in the control and conditional *Ift74* knockout mice by light microscopy.** Sperm from the control mice (a) showed normal appearance. Few sperm were recovered from the conditional *Ift74* knockout mice (b to d), and none of the sperm discovered showed normal morphology.

**Fig S3. Additional SEM images of epididymal sperm from the conditional *Ift74* knockout mice.** Images in “a” and “b” show two sperm with short tails and abnormal heads.

**Fig S4. Low magnification images of histology of testis and epididymis of adult control and conditional *Ift74* mutant mice.** A. Testis sections. Notice that seminiferous tubules in the control mice (upper) contain normally developed germ cells, and sperm were found in the lumen (arrow). In the mutant mice (lower), the seminiferous tubules were almost empty in the lumen. B. Epididymal sections. The epididymal lumens of a control mouse (upper) are filled with well-developed sperm. In *Ift74* mutant (lower), the cauda epididymal lumen contains sloughed spermatids, numerous detached sperm heads and abnormal tails and other cellular debris.

**Fig S5. Additional ultrastructural changes in the testis of conditional *Ift74* knockout mice.** a. The lumen of seminiferous tubule had degenerated cells and axoneme structure were hardly seen; b: misorganized microtubules (arrows) in the cytoplasm of an abnormally developed spermatid; c, d: abnormally condensed chromatin (arrowse: abnormally condensed chromatin (arrow), mis-organized mitochondria (dashed arrows); f, g: misorganized mictotubules (arrows); h: an abnormally localized axoneme (arrow), representing an abnormal spermatid that is being phagocyted by a Sertoli cell.

